# HBV vaccination and PMTCT as elimination tools in the presence of HIV: insights from a clinical cohort and dynamic model

**DOI:** 10.1101/162594

**Authors:** Anna L McNaughton, José Lourenço, Louise Hattingh, Emily Adland, Samantha Daniels, Anriette Van Zyl, Connie S Akiror, Susan Wareing, Katie Jeffery, M Azim Ansari, Paul Klenerman, Philip J R Goulder, Sunetra Gupta, Pieter Jooste, Philippa C Matthews

**Author notes:** These three authors contributed equally to the work presented here (Anna L McNaughton, José Lourenço, Louise Hattingh). Corresponding author Philippa Matthews; Address: Medawar Building for Pathogen Research, South Parks Road, Oxford OX1 3SY, UK.

## Abstract

**Background:** Sustainable Development Goals set a challenge for the elimination of hepatitis B virus (HBV) infection as a public health concern by the year 2030. Deployment of a robust prophylactic vaccine and enhanced interventions for prevention of mother to child transmission (PMTCT) are cornerstones of elimination strategy. However, in light of the estimated global burden of 290 million cases, enhanced efforts are required to underpin optimisation of public health strategy. Robust analysis of population epidemiology is particularly crucial for populations in Africa made vulnerable by HIV co-infection, poverty, stigma and poor access to prevention, diagnosis and treatment.

**Methods:** We here set out to evaluate the current and future role of HBV vaccination and PMTCT as tools for elimination. We first investigated the current impact of paediatric vaccination in a cohort of children with and without HIV infection in Kimberley, South Africa. Second, we used these data to inform a new model to simulate the ongoing impact of preventive interventions. By applying these two approaches in parallel, we are able to determine both the current impact of interventions, and the future projected outcome of ongoing preventive strategies over time.

**Results:** Existing efforts have been successful in reducing paediatric prevalence of HBV infection in this setting to <1%, demonstrating the success of the existing vaccine campaign. Our model predicts that, if consistently deployed, combination efforts of vaccination and PMTCT can significantly reduce population prevalence (HBsAg) by 2030, such that a major public health impact is possible even without achieving elimination. However, the prevalence of HBV e-antigen (HBeAg)-positive carriers will decline more slowly, representing a persistent population reservoir. We show that HIV co-infection significantly reduces titres of vaccine-mediated antibody, but has a relatively minor role in influencing the projected time to elimination. Our model can also be applied to other settings in order to predict time to elimination based on specific interventions.

**Conclusions:** Through extensive deployment of preventive strategies for HBV, significant positive public health impact is possible, although time to HBV elimination as a public health concern is likely to be substantially longer than that proposed by current goals.

## Background

The vaccine against hepatitis B virus (HBV) infection is one of the cornerstone strategies underpinning progress towards Sustainable Development Goals (SDGs) targets for HBV elimination as a public health threat by the year 2030 (1,2). Populations in southern Africa are particularly vulnerable to HBV-related morbidity and mortality due to the high prevalence of infection (≥8% in many regions) (3–5), co-endemic HIV infection (6), poor access to screening and diagnostics, limited access to antiviral therapy, stigma, and chronic neglect of education, research and resources (7,8). In this region, a substantial burden of HBV transmission occurs early in life, either vertically from mother to child, or through horizontal acquisition in young children (9). The HBV vaccine has been progressively rolled out as part of the World Health Organisation (WHO) Expanded Programme on Immunisation (EPI) over the past two decades (9), but the first vaccine dose is often postponed until age six weeks, when it is given together with other routine immunizations (10).

Vaccine deployment can be difficult to measure, as many children in Africa are born outside healthcare settings, there are no robust data regarding coverage of the three dose regimen (9), and different immunological correlates of protection have been applied (11,12). To accelerate progress towards elimination goals, suggested modifications to vaccine schedules have included shifting the first dose to be given at birth (13), additional doses in the context of HIV infection (14,15), booster doses in individuals whose antibody titre fails to meet a target threshold (11), and catch-up vaccination campaigns for adolescents and adults. However, there is a lack of robust data to inform which of these measures, individually or in combination, is most effective. Given the resource limitations of many settings in which HBV represents a public health challenge, there is an urgent need to underpin interventions with an evidence base derived both from careful observation of the existing impact of vaccination and from projections regarding ongoing impact.

On these grounds, we have set out to collect a detailed dataset to provide a snapshot of a population in South Africa in which HBV and HIV infections are co-endemic, first seeking evidence of the impact of the current immunization schedule in children, and then assessing the extent to which interventions could be predicted to achieve elimination targets. Finally, we built on this framework by adding data assimilated from the wider published literature to model the effects of different HBV vaccine deployment strategies, either alone or in combination with enhanced measures for prevention of mother to child transmission (PMTCT) through screening, antiviral therapy, accelerated immunisation and HBV immunoglobulin (16).

To date, few attempts have been made to model the impact of HBV vaccination, with one study modelling the global prevalence of current intervention efforts (17), and another that scrutinizes the combined impact of broad HBV elimination strategies (18). In this instance, we report a novel approach founded on primary clinical data, quantifying the individual and combined impact of childhood vaccination and PMTCT, and addressing the specific impact of co-endemic HIV infection. Combining output from a clinical dataset together with a dynamic model provides a synergistic approach to characterising the problem and projecting the effects of vaccination. Taking the evidence together, we conclude that while vaccination is a fundamental part of global elimination strategy and is highly effective in preventing infection in individual children, there remains an urgent need for rigorous, enhanced deployment of parallel strategies including education, diagnostics, antiviral therapy, and the ongoing quest for a cure.

## Methods

### Ethics Approval

Ethics approval was obtained from the Ethics Committee of the Faculty of Health Science, University of the Free State, Bloemfontein, South Africa (HIV Study Ref: ETOVS Nr 08/09 and COSAC Study Ref: ECUFS NR 80/2014). Written consent for enrollment into the study was obtained from the child’s parent/guardian.

### Study cohorts

Recruitment was undertaken in Kimberley, South Africa. In this setting, a standard three-dose HBV immunisation schedule is deployed in infants, with the first dose at six weeks. A previous study of HBV serology in adults in the same setting found HBsAg prevalence of 9.5% (55/579) (4). Children were recruited as part of the <u>C</u>o-infection in <u>S</u>outh-<u>A</u>frican <u>C</u>hildren (‘COSAC’) study as previously described (19,20). The lower age limit of recruitment was 6 months in order to limit the detection of maternal anti-HBs.

Children were recruited as follows:

1. HIV-negative children age 6-60 months (n=174), recruited through the Kimberley Respiratory Cohort (KReC) as previously described (19). These children were admitted to hospital between July 2014 and August 2016 with a clinical diagnosis of respiratory tract infection. KReC children were confirmed HIV-negative in 163 cases (93.7%). A further 11 children did not have an HIV test result recorded, but were assumed to be HIV-negative based on the clinical data recorded at the time of admission to hospital.
2. HIV-positive children were recruited primarily from HIV out-patient clinics between September 2009 and July 2016 as previously described (19,20). We recorded date of commencement of anti-retroviral therapy (ART), CD4+ T cell count and percentage, and HIV RNA viral load using the time point closest to the sample that was analysed for HBV serology. For the purpose of analysis, we divided these into two groups according by age:

i. Age 6-60 months; n=136. This group was selected to match the age range of the HIV-negative group, and also included five children who were initially screened for the KReC cohort but tested HIV-positive.
ii. Age >60 months (range 64-193 months); n=92.

At the time of undertaking this study, children were immunised with three doses of a monovalent HBV vaccine (Biovac Paed). Where possible, we recorded the number of HBV vaccine doses received based on the Road to Health Book (RTHB). The characteristics of the cohorts are summarised in table 2 and all metadata can be found in Suppl. data 1.

### Laboratory assessment of HBV status

Testing for Hepatitis B serum markers and DNA was performed as previously described, and in keeping with recent implementation of HBV screening in Kimberley (20). Briefly, HBsAg testing was carried out in Kimberley Hospital, South Africa using the Magnetic parcel chemiluminometric immunoassay (MPCI; Advia Centaur platform). Confirmatory HBsAg testing was carried out by the clinical microbiology laboratory at Oxford University Hospitals (OUH) NHS Foundation Trust, Oxford, UK (Architect i2000). For all samples, anti-HBs and anti-HBc testing were carried out by the OUH laboratory (Architect i2000). Limit of detection of the anti-HBs assay was 10 mIU/ml.

### Threshold for vaccine mediated immunity

Studies variably quote anti-HBs titres of ≥10 mIU/ml or ≥100 mIU/ml as a correlate of protection; UK recommendations for testing HBV immunity advocate the more stringent criterion of an anti-HBs titre of ≥100 mIU/ml (11), while early vaccine studies suggest a titre of ≥10 mIU/ml as a clinically relevant threshold for protection (12,21). We have presented our results pertaining to both thresholds.

### Statistical analysis

Data from the cohort was analysed using GraphPad Prism v.7.0. We determined significant differences between sub-sets within the cohort using Mann-Whitney U tests for non-parametric data, Fisher’s exact test for categorical variables and Spearman’s correlation coefficient for correlation between data points.

### Mathematical model of HBV transmission and prevention

We developed a dynamic model based on ordinary differential equations (ODE), for which parameterization of HBV transmission and prevention was based both on our Kimberley paediatric cohort and current literature estimates. This model was fitted to the HBV transmission background (Kimberley cohort) using Bayesian Markov Chain Monte-Carlo (bMCMC), and stochastic simulations were used to project impact interventions under various strategic scenarios. The full modelling and fitting approaches are detailed in Supplementary File 2.

### Measuring impact of interventions

SDGs for the year 2030 have been set out in the WHO Global Health Sector Strategy on Viral Hepatitis (GHSSVH) (2). Given the public health relevance of chronic infections, in particular of HBeAg-positive infections, we measured impact of interventions based on two targets:

i. The WHO target for a 90% reduction in HBsAg incidence, based on the assumption that this applies to chronic infection. (WHO goals also refer to HBsAg prevalence, and we present this in Supplementary File 2).
ii. An additional target for reduction of HBeAg-positive prevalence to 1 in 1000 (0.1%) in the whole population.

## Results

### Serological evidence of exposure to HBV infection

From our cohort of 402 children in Kimberley, South Africa, three were HBsAg-positive (0.7%; Table 1), indicating active infection. This HBsAg prevalence is significantly lower than in adults in a comparable study population (e.g. 11.1% in a previous study (4); p<0.0001). Exposure to HBV infection was measured using anti-HBc antibody; this was detected in three children (0.7%), one of whom was also HBsAg-positive. The other two were HBsAg-negative, indicating previous HBV exposure and clearance.

**Table 1.**
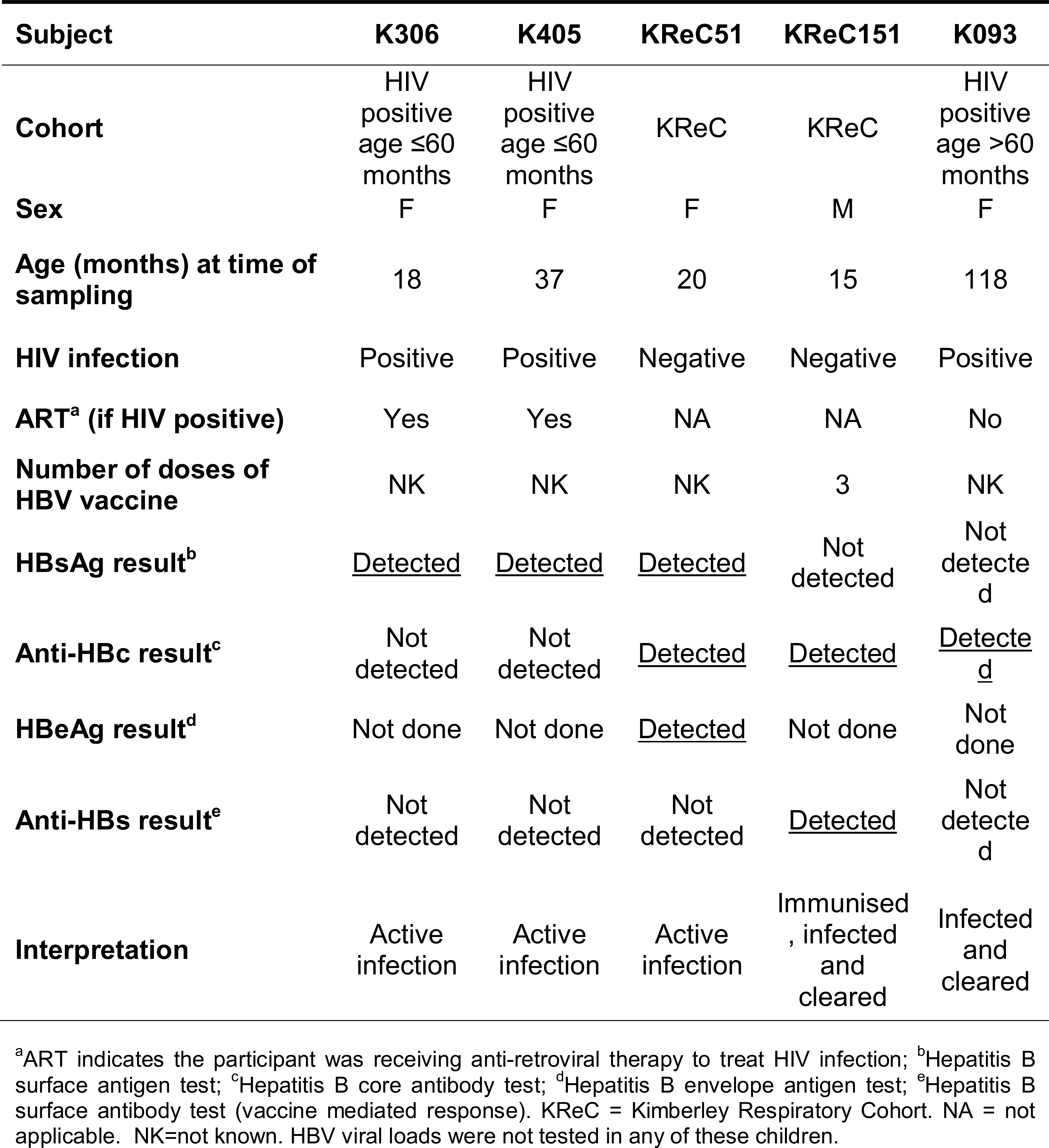
Profiles of five children from Kimberley, South Africa, with serological evidence of current or previous infection with HBV (based on positive HBsAg (n=3) or anti-HBc (n=2))

**Table 2.** Characteristics of three paediatric study cohorts, comprising 402 children, recruited from Kimberley Hospital, South Africa.

| **Cohort** | **HIV negative;**<br>**KReC**<br>**(age ≤60 months)** | **HIV positive**<br>**(age ≤60 months)** | **HIV positive**<br>**(age >60 months)** |
| --- | --- | --- | --- |
| **Number of subjects** | 174 | 136 | 92 |
| **Age range in months** | 8-58 | 6-60 | 64-193 |
| **Median age in**<br>**months (IQR)** | 18 (12-26) | 29 (18-40) | 137 (122-154) |
| **Sex (% male)** | 55.4 | 44.9 | 45.6 |

KReC = Kimberley Respiratory Cohort. IQR = interquartile range.

### Evidence of vaccination and immunity to HBV in children aged ≤ 60 months

We collected written evidence of immunisation from the RTHB in 90.8% HIV-negative (KReC) subjects and 6.3% of HIV-positive subjects (total 41.3% of cohort). None of the HBsAg-positive children attended with a written vaccination record. Among those with a RTHB record, 81.3% of HIV-negative and 100% of HIV-positive children were recorded as having received three HBV vaccine doses. Among all children age ≤60 months, 238/310 (77%) had an anti-HBs titre ≥10 mIU/ml suggesting some degree of vaccine-mediated immunity. The median anti-HBs titre in HIV-negative children was significantly higher than among the HIV-positive group (196 mIU/ml, vs. 11 mIU/ml, respectively, p<0.0001) (Fig 2A). There was no detectable anti-HBs antibody in 3.4% of HIV-negative vs. 47.8% of HIV-positive children (p<0.0001). Irrespective of the antibody titre used as a threshold for immunity, anti-HBs was higher in HIV-negative compared to HIV-positive children (Fig 2B). There was no significant difference in anti-HBs titres between male and female participants, either with or without HIV infection (p=0.49 and 0.31 respectively, data not shown).

**Figure 1.**
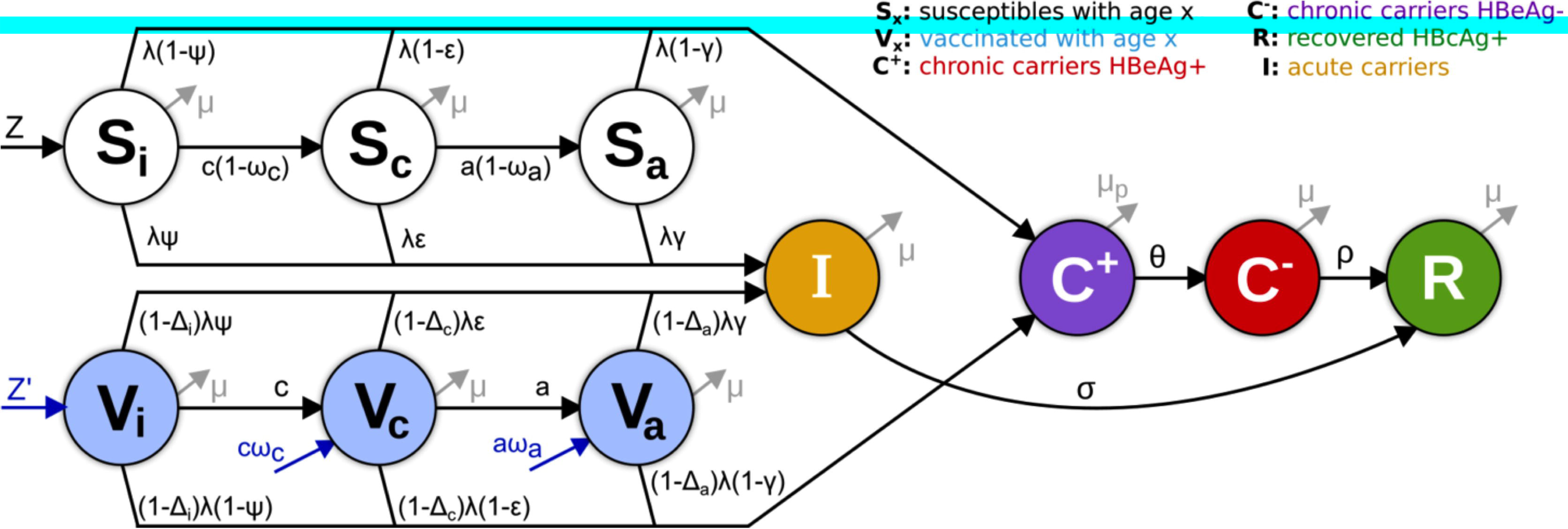
Diagram of HBV dynamic model. To allow for specific parameterization of important epidemiological states, the population was divided into susceptibles (S_x_) and vaccinated (V_x_) under 3 age-groups representing infants (x=*i*, <1 years of age), children (x=*c*, 1-6 years of age) and older individuals (x=*a*, >6 years of age). Individuals were assumed to acquire infection at any age, moving with different probabilities (Ψ, ε, γ, with Ψ < ε < γ) into acute (I) or chronic (C) infection. When chronically infected, individuals transit between HBeAg-positive (C+) and HBeAg-negative (C-) with rate θ, and may clear infection (R) with a small rate ρ. Vaccine-induced protection is age dependent (Δi) and assumed to lower susceptibility to infection (λ). Interventions (in blue) include routine vaccination at birth (Z’) and other ages (ωa, ωc), as well as PMTCT at birth (Z, Z’) and catch-up events (not shown). Model is used to fit prevalence rates as observed: HBV prevalence (I + C- + C+), HBcAb+ (R) and relative prevalence of HBeAg-positive (C+) and HBeAg-negative (C-) individuals. For a complete description on state transitions, vaccination, force of infection, parameters and model equations please refer to Suppl Data 2; within this document, Bayesian parameter estimations obtained when fitting the model are presented in Suppl Figure 1.

**Figure 2.**
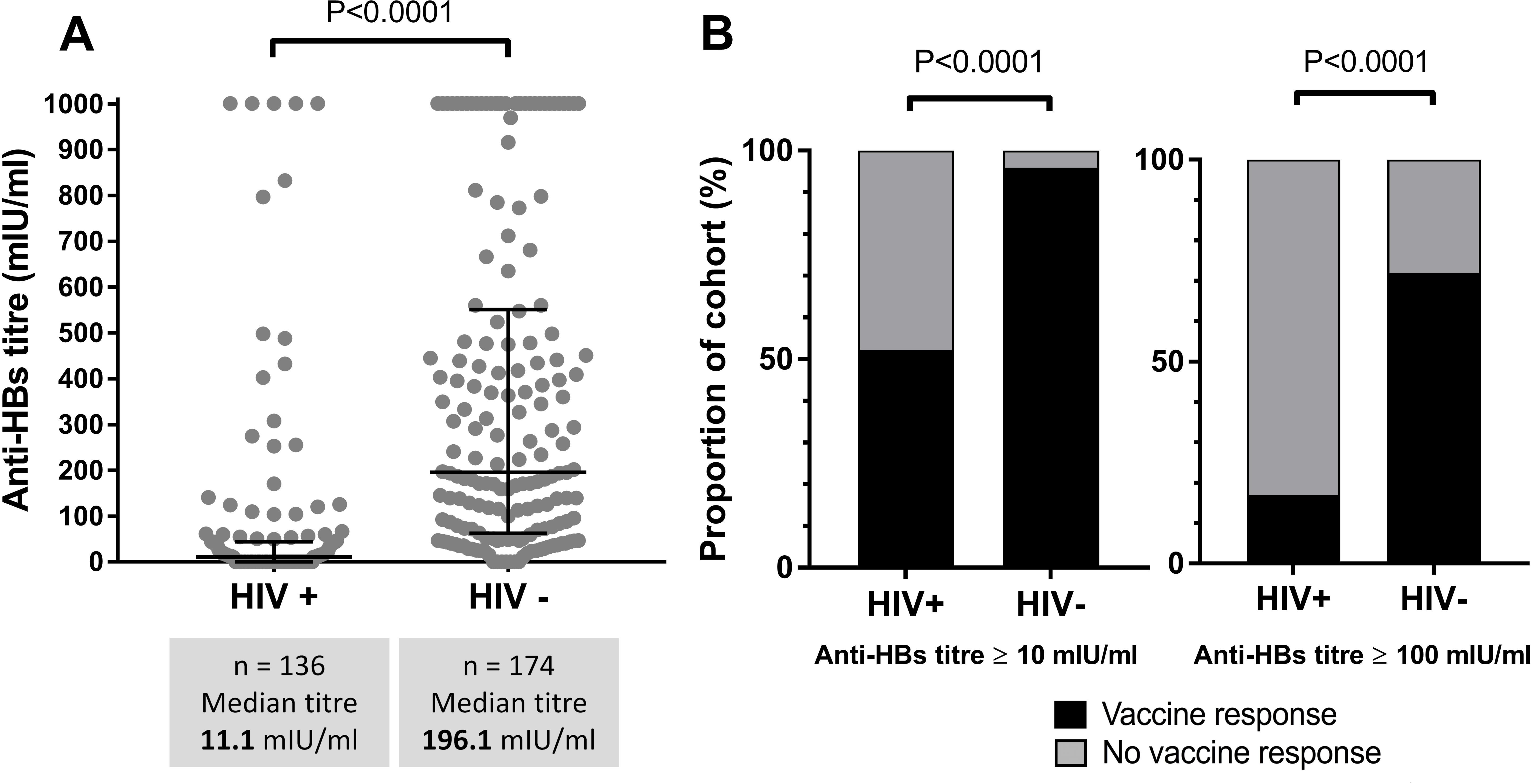
Hepatitis B surface antibody (anti-HBs) titres mediated by vaccination in HIV-positive (HIV+) and HIV-negative (HIV-) children aged 6-60 months in Kimberley, South Africa. A: Scatter plot representing vaccine-mediated antibody titres, indicating median and interquartile ranges (p-value by Mann Whitney U test). B: Proportion of children with anti-HBs ≥10 mIU/ml or ≥100 mIU/ml (p-values by Fisher’s Exact Test).

### Waning of vaccine response with age

HIV-positive children with anti-HBs titres ≥100mIU/ml were significantly younger than those with lower antibody titres (median age 17 months vs. 31 months, p=0.0008), while no such difference was observed within the HIV-negative group (Fig 3A). Using the lower threshold of ≥10mIU/ml, we found no significant difference by age in either the HIV-positive or the HIV-negative groups (p=0.17 and 4.48 respectively, data not shown). To expand our view of the HIV-positive group, we also added analysis of an older cohort (92 children aged >60 months), and demonstrated that anti-HBs titres were significantly lower in this older group (p<0.0001), with only 2/92 subjects (2.2%) achieving a detectable anti-HBs titre (Fig 3B). Anti-HBs titres waned significantly with age up to age 60 months in HIV-positive children (Fig 3C; p=0.004). We observed a similar trend in the HIV-negative cohort, but this did not reach statistical significance (Fig 3C; p=0.07). The proportion of HIV-positive subjects with a detectable anti-HBs titre declined steadily with age in the cohort, contrasting to the trend in HIV-negative subjects, where individuals maintained protective anti-HBs titres despite a trend towards decreasing mean titres (Fig 3C).

**Figure 3.**
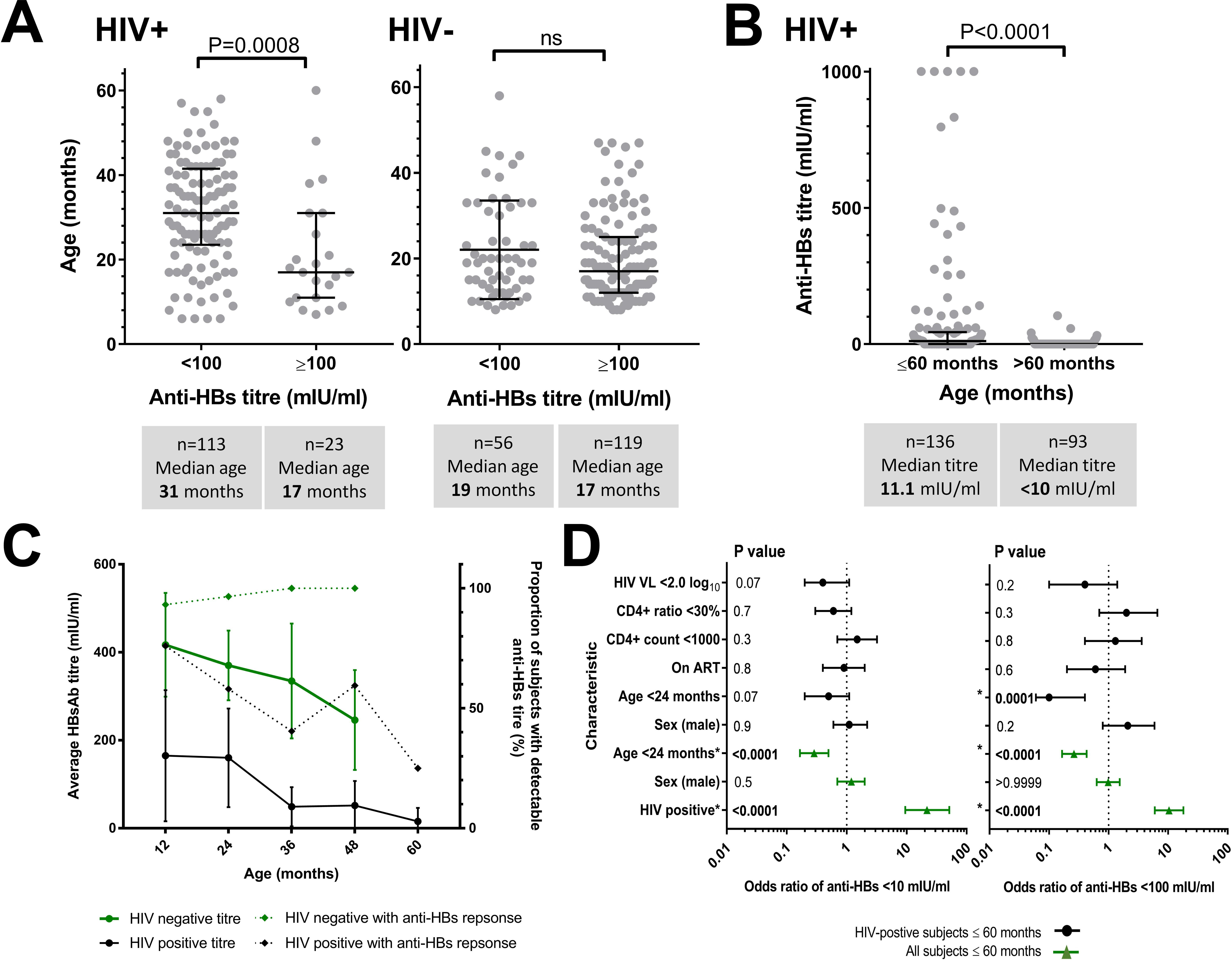
Relationship between age and vaccine-mediated Hepatitis B surface antibody (anti-HBs) titres in HIV-positive and HIV-negative children in Kimberley, South Africa. A: Ages of children attaining anti-HBs titres ≥100 mIU/ml for HIV-positive and HIV-negative children age 6-60 months. Median ages, interquartile ranges and p-values by Mann-Whitney U test are indicated. B: Relationship between age and vaccine-mediated Ab titre among HIV-positive children including those age 6-60 months and an older cohort age >60 months (range 64-193 months). P-value by Mann Whitney U test. C: Anti-HBs titre and proportion of subjects with a detectable titre for HIV-positive and HIV-negative children according to age. On the solid lines, each point represents the mean titre (with 95% confidence intervals) for the group of children aged ≤12 months (1 yr), 13-24 months (2 yrs), 25-36 months (3 yrs), 37-48 months (4 yrs), 49-60 months (5 years). For the same groups of children, the dotted lines represent the proportion of subjects with a detectable titre and the 95% confidence intervals. Trends within the data were assessed using linear regression analysis. D: Odds ratios for protective response to HBV vaccination in children age 6-60 months in Kimberley, South Africa are shown for anti-HBs titre <10mIU/ml and <100mIU/ml in the whole cohort (green) and in HIV-positive children (black). Statistically significant OR are denoted * and significant p-values are indicated in bold. Figure 2: Relationship between age and vaccine-mediated Hepatitis B surface antibody (anti-HBs) titres in HIV-positive and HIV-negative children in Kimberley, South Africa.

### Stratification of vaccine responses by anti-retroviral therapy (ART) among HIV-positive children

For HIV-positive children aged ≤60 months, ART treatment data were available for 79% of subjects. Within this group, 71% were receiving ART at the time we tested for anti-HBs, and had received a median of 20 months of treatment (IQR 6-33 months). Comparing anti-HBs titres between ART-treated vs. untreated children, we found no significant difference (p=0.72; 76 ART-treated, median anti-HBs 13.3 mIU/ml and 31 untreated children, median anti-HBs 14.1 mIU/ml, data not shown). There was also no difference between anti-HBs titres of children treated for ≤12 months vs. >12 months (p=0.50, data not shown). We did not examine the effect of ART on anti-HBs titres in children >60 months due to the low numbers of subjects with a detectable anti-HBs titre (n=2).

### Odds of developing an anti-HBs response

We used an odds ratio (OR) analysis to identify factors associated with vaccine-mediated protection (Fig 3D). HIV-positive status was associated with lack of protection, for antibody titres of both <10 mIU/ml (OR 26.2, 95% CI 11.2-58.6), and <100 mIU/ml(OR 11.6, 95% CI 6.7-20.4). In contrast, younger age(<24 months) was protective in HIV-positive subjects, (for anti-HBs <10 mIU/ml OR 0.3, 95% CI 0.2-0.5 and for anti-HBs <100 mIU/ml OR 0.3, 95% CI 0.2-0.4). Gender, ART, CD4+ count, CD4+ ratio and HIV viral load were not found to be significantly predictive of anti-HBs titres at either threshold.

### Fitting of the dynamic model to local HBV epidemiology

The model closely reproduced the fitted variables: HBV prevalence (HBsAg), prevalence of HBV exposure (anti-HBc) and relative proportion of HBeAg-negative and HBeAg-positive among chronic carriers (Suppl Fig 1 A12). Parameters for which insufficient support was found in the literature, the resulting posteriors were well behaved (Suppl Fig 1B), while for parameters using informative priors the posteriors matched well (Suppl Fig 1CD). Overall, the obtained bell-shaped posteriors highlighted a good data fit and no identifiability issues with the bMCMC approach. Based on these results, we next simulated stochastic projections of intervention impact.

### Model projection of the impact of routine neonatal vaccination and PMTCT alone

Based on SDGs (2), Fig 4 shows the projected impact of varying coverage of neonatal vaccination and PMTCT. Both HBsAg incidence (Fig 4 A1) and HBeAg-positive prevalence (Fig 4 B1) reduce faster with increasing neonatal immunization coverage, resulting in shorter times to reach SDGs (Fig 4 A2, B2). Importantly, even immunization of 100% of neonates is predicted to take 99 years (95% CI 61 - 186) for the HBsAg incidence target (Fig 4 A2), and ~175 years (95% CI 103 - 278) for the HBeAg prevalence target (Fig 4 B2).

**Figure 4.**
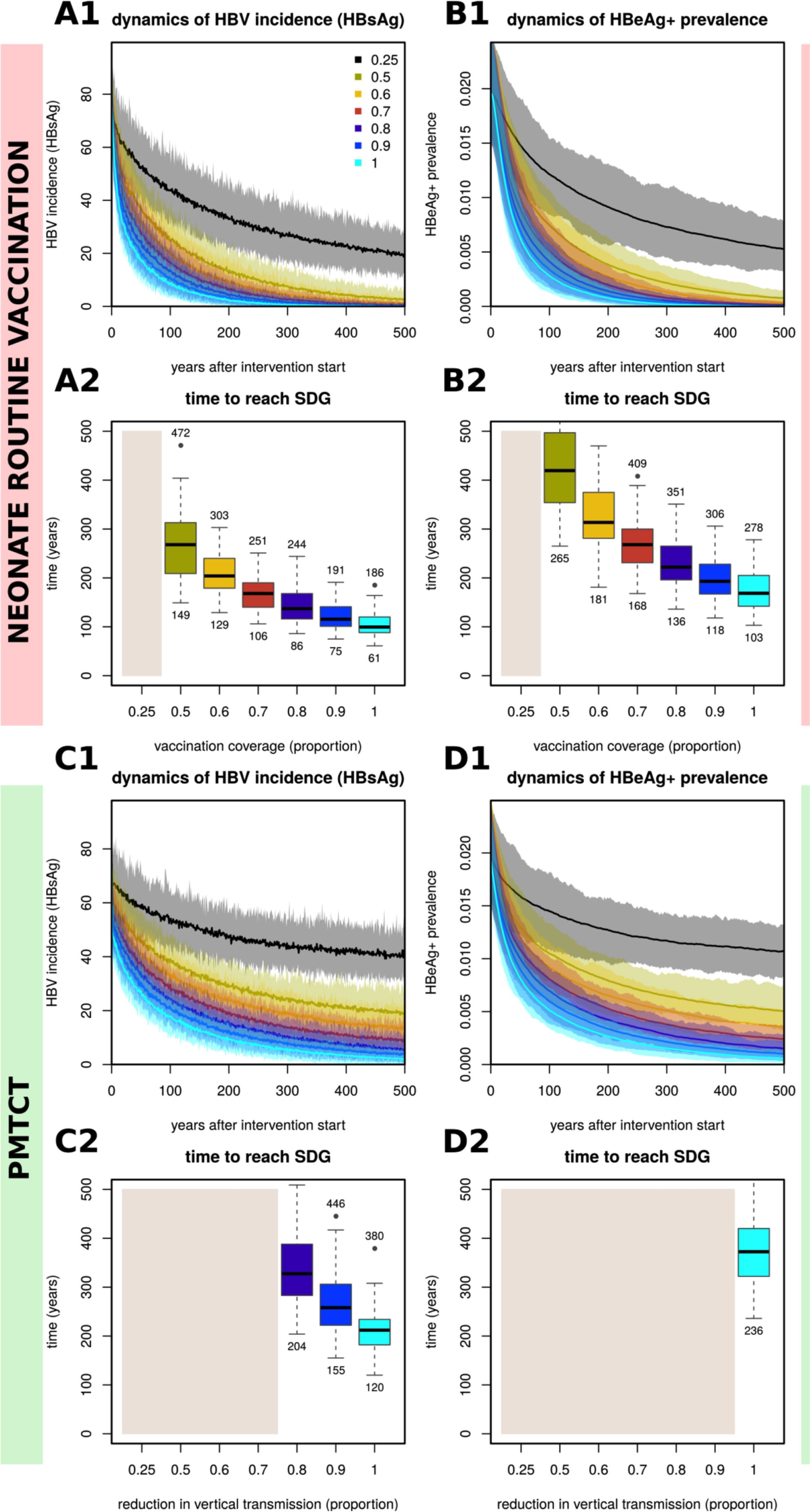
Stochastic impact of neonatal vaccination and PMTCT on HBV incidence (HBsAg) and HBeAg+ prevalence, showing time to reach sustainable development goals when using interventions independently. **(A1-A2)**Impact on **HBV incidence (HBsAg)** (A1) and time to reach sustainable development goal (SDG) (A2) for varying routine immunization coverage of neonates. **(B1-B2)** Impact on HBeAg+ prevalence (B1) and time to reach SDG (B2) for varying routine immunization coverage of neonates. **(C1-C2)** Impact on **HBV incidence (HBsAg)** (C1) and time to SDG (C2) for varying PMTCT coverage. **(D1-D2)** Impact on HBeAg+ prevalence (D1) and time to reach SDG (D2) for varying PMTCT coverage. **(A1, B1, C1, D1)** Lines are the mean and shaded areas the standard deviation of model output when running 50 stochastic simulations per intervention (sampling the parameter posteriors shown in Figure 1). **(A2, B2, C2, D2) HBV incidence (HBsAg)** SDG is set to a reduction of 90%. HBeAg+ prevalence SDG is set to 1/1000 individuals. Beige areas mark interventions reaching SDGs after 500 years on average. Boxplots show the variation of the 50 stochastic simulations. Numbers above and below boxplots show the 2.5% lower and 97.5% upper limits of the solutions. **(All suplots)** Intervention coverage varies from 0.25 to 1 (as coloured and named in subplot A1).

With PMTCT interventions, both HBsAg incidence (Fig 4 C1) and HBeAg prevalence (Fig 4 D1) reduced faster in time for increasing efforts, resulting in shorter times to reach the elimination targets (Fig 4 C2, D2). However, the impact of PMTCT was smaller than neonatal vaccination, resulting in longer times to reach SDGs. In fact, for the majority of PMTCT effort levels simulated, SDGs could not be reached within 500 years (beige areas in Fig 4 C2, D2). For HBeAg-positive prevalence, only when PMTCT effort was 1 (i.e. no vertical transmission), was the SDG attainable within 500 years.

For complementary results using neonatal vaccination and PMTCT with impact on total prevalence (acute and chronic) see Support Figure 4.

### Modelling progress towards HBV elimination by the year 2030 based on combinations of neonatal vaccination and PMTCT

We projected impact of combined interventions by the year 2030 (Fig 5 A1, B1), and predicted the year at which SDGs would be reached (Fig 5 A2, B2). Strikingly, HBsAg incidence could already have been reduced by >90% (Fig 5 A1) if both neonatal vaccination and PMTCT had been deployed at 100% coverage since they became available in 1995 (mean predicted year of elimination 2017; Fig 5 A2). In reality, complete coverage is not possible, and we therefore projected outcomes based on <100% coverage. For example, combining neonatal vaccination and PMTCT with 90% coverage of each since 1995 would achieve the HBsAg incidence target by 2028; if this is reduced to 80% coverage then goals would be attained by 2044. To achieve the target reduction in HBeAg prevalence, modelled on 90% coverage and 80% coverage of interventions, the projected years are 2072 and 2096 respectively (Fig 5 B1, B2).

**Figure 5.**
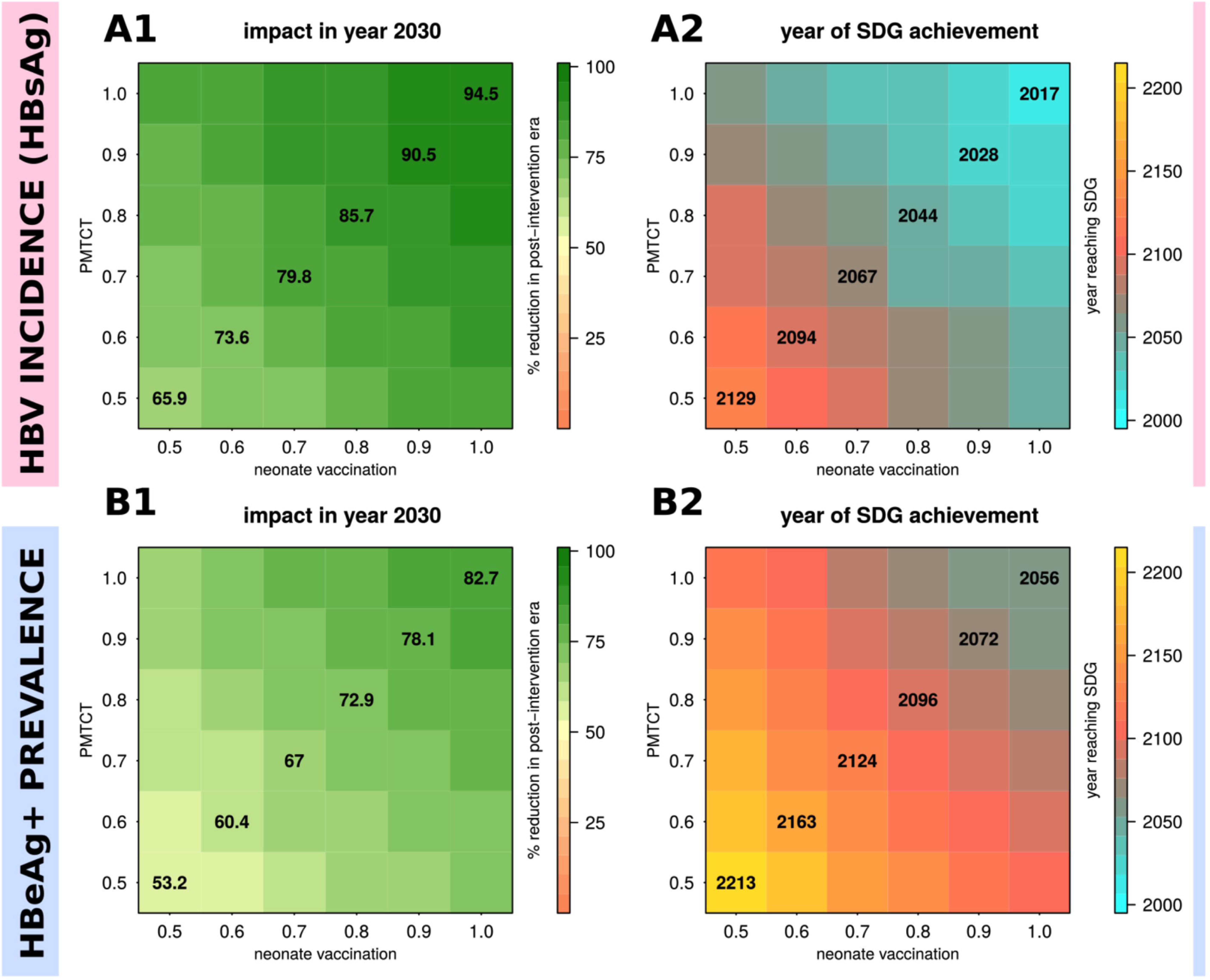
Sensitivity of mean intervention impact on HBV incidence (HBsAg) and HBeAg+ prevalence based on combinations of routine neonatal vaccination and PMTCT. **(A1-A2)** Mean impact of interventions on **HBV incidence (HBsAg)** (A1) and mean time to reach sustainable development goals (SDGs) (A2). **(B1-B2)** Mean impact of interventions on HBeAg+ prevalence (B2) and mean time to reach SDG (B2). **(All subplots)** Impact is shown as percent reduction in incidence or prevalence compared to pre-intervention levels (e.g. 50 indicates a 50% reduction compared to before the start of the intervention). **HBV incidence (HBsAg)** SDG is set to a reduction of 90%. HBeAg+ prevalence SDG is set to 1/1000 individuals. Mean results are obtained from 50 stochastic simulations per intervention combination (vaccination, PMTCT) with parameters sampled from the posteriors shown in Support Figure 1. Start of interventions in the stochastic simulations is in year 1995 to simulate an appropriate time scale to address impact by 2030.

Overall, the highest probability of achieving elimination targets is through a combination of 100% neonatal vaccination coverage and PMTCT (Fig 6 A1, red line). Based on 90% coverage of neonatal vaccine and PMTCT (Fig 6 A1, A2, green line) as proposed in WHO’S GHSSVH (2), there was only 50% probability of reaching the HBsAg incidence target by 2030, and approaching 100% probability only by 2050. For the target based on HBeAg prevalence, the probabilities of achieving the goal were pushed forward by approximately four decades.

**Figure 6.**
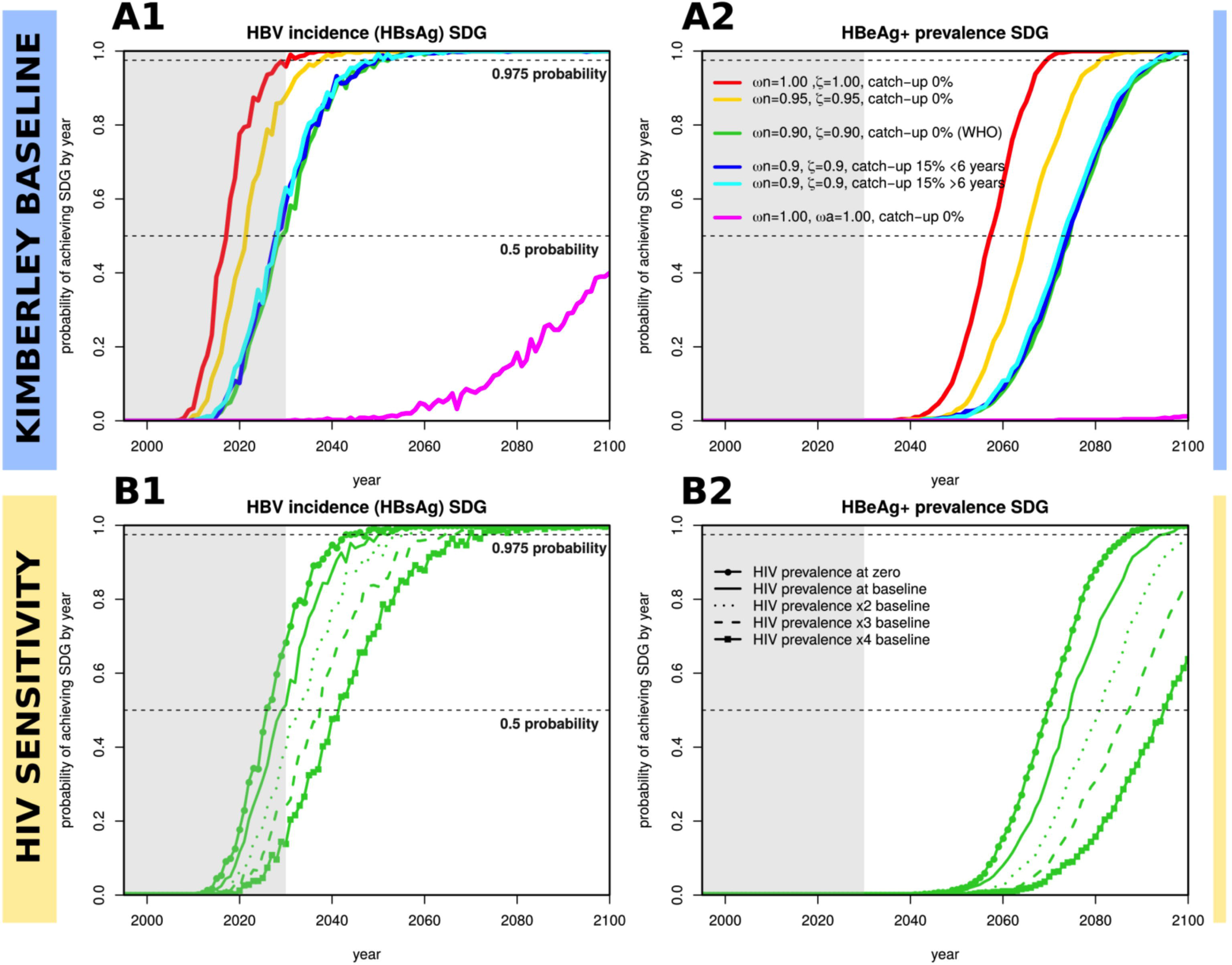
Yearly estimated probabilities of achieving sustainable development goals for HBV incidence (HBsAg) and HBeAg+ prevalence based on particular combinations of interventions and local HIV prevalence levels. A total of 1000 stochastic simulations are run independently for each set of particular interventions (coloured legend, subplot A2), with each using a random parameter sample from the posteriors shown in Support Figure 1. Interventions start in year 1995. For every year post-intervention start, the proportion of simulations that have achieved the sustainable development goals (SDGs) is recorded and taken to be the probability. **(A1)** Probability of reaching **HBV incidence (HBsAg)** SDG in time (goal is set to a reduction of 90%). **(A2)** Probability of reaching HBeAg+ prevalence SDG in time (goal is set to 1/1000 individuals). **(B1, B2)** Same as subplots A1-A2 but addressing sensitivity to HIV prevalence levels in the population for a particular intervention (green, ω*n=0.9*, ζ=0.9, catch-up 0% (WHO)). Solid line is the same as in subplots A1-A2 (named HIV prevalence at baseline). Other lines present results assuming zero HIV prevalence (full line with points) or higher prevalences (dotted, dashed, line with squares). **(All subplots)** The dashed horizontal lines mark 0.5 and 0.975 probability of achieving SDGs. The grey shaded area marks the time period before 2030. In the interventions, ω*n* is routine vaccination of neonates, ζ the PMTCT effort, ω*a* routine vaccination of +6 years of age, and catch-up a one-off event of vaccination in some age groups or general population.

For complementary results on impact and time to reach SDGs when considering combinations of PMTCT and routine vaccination at the age of 6 see Support Figure 5, and for combinations of PMTCT and neonate routine vaccination plus a complete catch-up campaign see Support Figure 6.

### Projecting the probability of achieving elimination targets based on combinations of neonatal vaccination, PMTCT and enhanced vaccination

We simulated the impact of combining neonatal vaccination and PMTCT with additional vaccine deployment (Fig 6 A1, A2), through the routine vaccination of older children (≥6 years of age), and one-off catch-up vaccination of children (<6 years) and others (>6 years). Adding catch-up vaccination campaigns makes no impact on the probability of reaching SDGs (Fig 6 A1, A2, blue and cyan lines). Routine vaccination at 6 years of age, even when delivered at 100% coverage, is markedly less effective than any other projected intervention (Fig 6 A1, A2, magenta line).

### Projecting the impact of HIV on the probability of achieving elimination targets

A baseline scenario was defined by the epidemiological setting fitted by our model in the context of Kimberley, using local HIV prevalence for each of the modelled age groups (Fig 6 B1, B2, solid line). In a sensitivity exercise, alternative scenarios were considered in which baseline HIV prevalence was altered to zero or higher prevalence. When compared with no HIV (Fig 6 B1, B2, dotted line), the presence of HIV at the prevalence seen in Kimberley (Fig 6 B1, B2, solid line) adds an estimated four years to achieve a 50% chance of reaching the goals (Fig 6 B1). Higher baseline HIV prevalence (x2, x3 and x4 baseline data for Kimberley) was used to investigate the potential impact of coinfection in high-risk populations. Thus, increasing HIV prevalence has a negative impact on the success of interventions for HBV, but the effects are relatively modest. In particular, doubling HIV prevalence would shift the 50% probability endpoint into the future by ~4 years for the HBsAg incidence target, and ~7 years for the HBeAg prevalence target.

## Discussion

This is a unique study in which we capitalize on detailed clinical cohort data collected in South Africa in order to form a robust view of the nature of vaccine-mediated immunity, and develop a mathematical model of HBV transmission and prevention. Overall, we demonstrate that the optimum population intervention is high coverage neonatal vaccination, and that this can be strengthened by robust deployment of PMTCT. However, we project long time-scales to achieve elimination targets, congruent with the large established reservoir of chronic HBV infection, lack of curative therapy, infection that can persist for the entire life-span of the host, and interventions that target only a small proportion of the population. Although a high coverage of neonatal vaccination combined with robust PMCTC shows potential promise, the projected time-frame for elimination is currently substantially beyond the 2030 milestone. Complete extinction of infection is far beyond reach based on currently available interventions, and current efforts should be focused on control of HBV as public health issue rather than complete elimination.

By assimilating the results of a clinical cohort study and a model, we develop a more complete picture than either individual approach would provide in isolation. Only by viewing the two conclusions together can we correctly infer that vaccination is of profound importance in protecting individual children and significantly reducing the burden of infection in paediatric cohorts, but also that continuing to pursue this strategy alone is not sufficient to bring about HBV elimination, or even robust control, within the desired time-scale.

Compared to published models of other vaccine-preventable diseases (22), there is a marked deficit in the existing literature for HBV, with few other modelling efforts represented in the literature (23,24). Reassuringly, our findings are consistent with those of another recent simulation of HBV prevention (18); we concur in concluding that current vaccine-based interventions will result in a modest reduction in HBV prevalence by the year 2030. However, there are also some important differences that distinguish our work from previous efforts:

i. Our evaluation provides the advantages of both clinical data and a mathematical model, with close links between our cohort and simulations, and strengths in interpretation of data derived through different approaches. In so doing, we have also been able to specifically address the impact of co-endemic HIV.
ii. We focus on a particular population for which we derive unknown epidemiological parameters and apply a robust data-driven approach to others. Our Bayesian framework therefore stands alone (as a tool) that can be applied to any population for which empirical support of key HBV epidemiological parameters is missing. By supplying the model’s code, we can facilitate the use of the tool by others.
iii. As outputs, we have used targets for reductions in both HBsAg incidence and HBeAg-positive prevalence, and have projected the impact of interventions based specifically on the WHO proposal for 90% vaccination of neonates and 90% PMTCT coverage by 2030. Previous studies (17,18) have focused instead on *ad hoc* control thresholds or impact on the public health problem through reduction of HBV-related deaths. Our results thus contribute to an ongoing discussion regarding which goals should be set, and their underlying public health implications. The model suggests that reaching either of the elimination targets will require different intervention coverage and different time scales. In particular, the target for reducing HBsAg incidence is easier to achieve than reducing HBeAg-prevalence.
iv. This is an important parsimonious, data-driven tool, offering the potential to scrutinise different strategies independently from one another.

### Impact of HIV on population interventions for HBV

Our results support previous evidence that HBV vaccine-mediated immunity wanes over time independently of HIV serostatus, but faster for HIV-positive individuals (25). Impaired vaccine responses have previously been reported in HIV-positive individuals (15,26–29), but it is also possible that vaccine coverage is lower in HIV-infected children (30). However, waning of anti-HBs titres does not necessarily correlate with loss of clinical protection; anamnestic responses are thought to occur in a proportion of those vaccinated (31), although this memory may be attenuated by HIV (32,33).

ART has previously been associated with improved HBV vaccine responses (34,35), although we did not replicate this finding in our cohort. This can potentially be explained by a previous study in the same setting, demonstrating that immune reconstitution takes a median of five years after ART initiation (36). Our current study is underpowered to detect any ART effect, given both the relatively short durations of therapy, and the small number of untreated children. Interestingly, despite the lack of direct association with ART, children with lower HIV viral loads had significantly higher anti-HBs titres, in keeping with previous studies (14,34).

Our cohort highlights day-to-day challenges of drug provision and monitoring in this setting: we did not have access to detailed prospective ART treatment data, guidelines have changed numerous times since 2002, and lamivudine (3TC) was intermittently used as a substitute for nevirapine (NVP) due to supply issues. During the period covered by our study, ART was only introduced in children achieving certain immunological criteria, while new guidelines recommend that all HIV-infected children are started on ART (37). The immune reconstitution of this population over time is likely to reduce differences between HIV-positive and HIV-negative groups. ART treatment is relevant to outcomes in individuals with HIV/HBV coinfection, as first line ART regimens include either 3TC or tenofovir (TDF), both of which have activity against HBV. Alternative approaches for HBV prevention in HIV-positive subjects, such as supplementing the current schedule with booster vaccinations and increased vaccine doses have been trialled with variable results (14)(15).

Our projections propose that HIV does have a negative effect on HBV interventions, although HIV prevalence only marginally increases time to reach elimination targets, which may not be significant in light of the long overall time-frames projected, even in the absence of HIV. The high HIV prevalences modelled can occur in specific high-risk groups including sex workers and men who have sex with men (38) and it is likely that increased intervention will be required in these groups to minimise HBV transmission.

### Changes required to meet international goals

The model suggests long time-lines, enumerated in centuries rather than decades, before control targets are reached using vaccination or PMTCT alone. Combinations of these interventions show much shorter time scales. Based on currently available interventions, major scaling up of both neonatal vaccination and PMTCT efforts will be required. Importantly, the prevalence of HBeAg-positive carriers, who are at an elevated risk of chronic liver disease and hepatocellular carcinoma, as well as being at higher risk of transmitting their infection, will decline at a slower rate. Setting a control target based on reduction in the number of new HBV cases (i.e. HBsAg incidence) can therefore lead to the most optimistic projections but distract attention from the importance of reducing HBeAg prevalence.

Our results also underscore that a major public health impact is possible, even without achieving elimination. Careful adjusting of expectations and aims, according to the scale on which particular changes occur, may inform the setting of realistic targets (e.g. reduction in the prevalence of HBeAg could be the most informative outcome measure). The wrong choice of either target or timescale may result in unnecessary abandonment of a strategy that could have a major impact in a few decades. In addition to informing rational use of interventions that have a positive population impact, our study is also important in cautioning against the use of strategies that may have little or no lasting population impact. This is illustrated by our results for catch-up HBV vaccination, which adds little in situations where high coverage of both neonatal immunization and PMTCT can be attained.

### Caveats and limitations

Different approaches to recruitment of our HIV-positive and HIV-negative cohorts may have introduced unintentional bias. The KReC children may be less healthy than HIV-negative children in the community, and this approach to recruitment predominantly selected younger children (on average 9.4 months younger than the HIV-positive cohort).

We set out to focus on children aged <60 months in order to collect data from the RTHB. However, in practice, we did not capture good RTHB data and data collection from the RTHB is itself subject to bias, as families who attend with such records may be those who are most likely to have immunised their children. Numerous complex social factors are also relevant in determining immunisation status; babies born to mothers who have HIV and/or HBV are more likely to be in disadvantaged by poverty, and by illness and death in the family, such that they might be less likely to present for (or respond to) vaccination. However, in this setting (and others where antenatal HBV screening is not routinely deployed (7,39,40)), we deem it unlikely that there is a significant difference in vaccination rates between infants born to HBV-positive versus HBV-negative mothers.

HBV DNA is a more sensitive screening tool than HBsAg but was not practical due to high cost and lack of availability in this setting. The relatively small numbers in each age group and the lack of longitudinal follow-up for individual children puts limitations on the data showing anti-HBs waning over time, but the trends we observe here are biologically plausible and consistent with existing literature (25,41).

Although we have estimated and parameterized the impact of HIV status on HBV vaccine-induced protection, we have not modelled other factors related to HIV infection. Namely, we have not included the potential for increased susceptibility to HBV infection or increased risk of vertical transmission. These factors would have required further model classes and specific parameterization, for which little literature support exists. We have also not considered the influence of population migration on the success of HBV interventions to reach the elimination targets. Migration of non-immune and/or infected individuals into an area would delay the time to achieve the targets estimated by our modelling approach.

## Conclusions

Our results affirm the success of the HBV vaccine programme in reducing the prevalence of HBV in children, with current paediatric prevalence rates of <1%. However, we also highlight that cases of HBV transmission persist and that a proportion of children are potentially at risk of infection as a result of low anti-HBs titres, either as a result of missing or incomplete immunisation, or because of poor antibody titres following vaccination. We predict that current elimination targets, in particular when framed around reductions of HBeAg-positive prevalence, are unlikely to be achieved by 2030. For optimum impact, we suggest that elimination targets should be defined around HBeAg-positive carriers, which are a major proxy for the public health burden of HBV. Our study highlights the essential need to collect better data that can help to inform progress towards targets, to optimize deployment of vaccination and PMTCT, and to invest substantially in education, case finding and treatment. The prospects of control would be substantially enhanced by improvements in therapy, and ultimately, the only route to elimination of HBV may be to develop a cure.

## Declarations

### Ethics approval and consent to participate

Ethics approval was obtained from the Ethics Committee of the Faculty of Health Science, University of the Free State, Bloemfontein, South Africa (HIV Study Ref: ETOVS Nr 08/09 and COSAC Study Ref: ECUFS NR 80/2014). Written consent for enrollment into the study was obtained from the child’s parent/guardian.

### Consent for publication

Not applicable

### Availability of data and materials

The datasets generated and/or analysed during the current study are available in the Figshare repository: https://figshare.com/s/cd1e4f324606949d1680 Model code will be made available on GitHub on acceptance of the article for publication.

### Competing interests

PCM is an Associate Editor for BMC Infectious Diseases and undertakes consultancy work for Immunocore.

### Funding

PCM, PK and PJRG are funded by the Wellcome Trust (grant numbers 110110/Z/15/Z to PM, 109965MA to PK, and 104748MA to PJRG); https://wellcome.ac.uk. Recruitment and serological testing of the KReC cohort was covered by a project grant awarded to PCM from the Rosetrees Trust http://www.rosetreestrust.co.uk/. SG and JL received funding from the European Research Council under the European Union’s Seventh Framework Programme (FP7/2007-2013)/ERC grant agreement no. 268904-DIVERSITY https://erc.europa.eu/. PK is also funded by an NIHR Senior Fellowship https://www.nihr.ac.uk/. The funders had no role in study design, data collection and analysis, decision to publish, or preparation of the manuscript.

### Authors’ Contributions

- Conceived the idea: PM, SG, PJ
- Applied for ethical permission: PM, PG, PJ
- Secured funding: PM, PG, PJ, PK
- Wrote the manuscript draft: AM, JL, PM
- Recruited children and collected clinical data: LH, EA, SD, AvZ, PJ
- Analysed clinical data: EA, CA, AM
- Undertook laboratory assays for HBV serology: SW, KJ
- Developed and tested mathematical model: JL, SG with input from AM, PM, MAA
- All authors read and approved the final manuscript.

## Acknowledgements

Not applicable

## Supplementary Files

**Suppl data 1 – Cohort metadata.** Metadata for three paediatric cohorts recruited in Kimberley, South Africa, including longitudinal CD4+ T cell and viral load data for paediatric HIV cohort age ≤60 months in Kimberley, South Africa. This file is available on-line via the following link: https://figshare.com/s/cd1e4f324606949d1680.

Suppl data 2 – Mathematical model and supporting figures.Complete model description, including parameters, assumptions, mathematical expressions and fitting details, further including results and figures supporting the main text.

## Abbreviations

HBV: Hepatitis B virus
SDGs: Sustainable Development Goals
HIV: Human Immunodeficiency Virus (type 1)
WHO: World Health Organisation
EPI: Expanded Programme on Immunisation
PMTCT: Prevention of mother to child transmission
COSAC: Coinfection in South African children
HBsAg: Hepatitis B surface antigen
Anti-HBs: Antibody to hepatitis B surface antigen (vaccine-mediated antibody)
KReC: Kimberley Respiratory Cohort
RTHB: Road to Health Book
Anti-HBc: Antibody to hepatitis B core antigen (antibody mediated by exposure to infection)
ODE: Ordinary differential equation
HBeAg: Hepatitis B envelope antigen
GHSSVH: Global Health Sector Strategy on Viral Hepatitis
ART: Anti-retroviral therapy
3TC: Lamivudine
OUH: Oxford University Hospitals

